# Competitive inter-species interactions underlie the increased antimicrobial tolerance in multispecies brewery biofilms

**DOI:** 10.1101/204628

**Authors:** Ilse Parijs, Hans P. Steenackers

**Affiliations:** Centre of Microbial and Plant Genetics (CMPG), Department of Microbial and Molecular Systems, KU Leuven, Kasteelpark Arenberg 20 - box 2460, B-3001 Leuven, Belgium

## Abstract

Genetic diversity often enhances the tolerance of microbial communities against antimicrobial treatment. However the sociobiology underlying this antimicrobial tolerance remains largely unexplored. Here we analyze how inter-species interactions can increase antimicrobial tolerance. We apply our approach to 17 industrially-relevant multispecies biofilm models, based on species isolated from 58 contaminating biofilms in three breweries. Sulfathiazole is used as antimicrobial agent because it shows the highest activity out of 22 biofilm inhibitors tested. Our analysis reveals that competitive interactions dominate among species within brewery biofilms. We show that antimicrobial treatment can reduce the level of competition and therefore cause a subset of species to bloom. The result is a lower percentage inhibition of these species and increased tolerance. In addition, we show that the presence of competing species can also directly enhance the inherent tolerance of microbes to antimicrobial treatment, either because species protect each other or because they induce specific tolerance phenotypes as a response to competitors (i.e. competition sensing). Overall, our study emphasizes that the dominance of competitive interactions is central to the enhanced antimicrobial tolerance of the multispecies biofilms and that the activity of antimicrobials against multispecies biofilms cannot be predicted based on their effect against mono-cultures.

## INTRODUCTION

Microbes commonly live in surface-attached communities embedded in a self-produced matrix, known as biofilms, which cause major problems and economic losses within industrial and medical sectors (Hall-Stoodley *et al.*, 2004). The majority of natural biofilms contain multiple species and harbor different functions and abilities compared to their monospecies counterparts (Stoodley *et al.*, 2002; Elias and Banin, 2012). One hallmark of multispecies biofilms is their increased tolerance against antimicrobial agents (Baffone *et al.*, 2011; Simões *et al.*, 2010; Shakeri *et al.*, 2007; Kumar and Peng, 2015; Jagmann *et al.*, 2015; Adam *et al.*, 2002; Lopes *et al.*, 2012; Leriche *et al.*, 2003; Whiteley *et al.*, 2001; Luppens *et al.*, 2008; Wang *et al.*, 2013; Schwering *et al.*, 2013; Van der Veen and Abee, 2011; Simões *et al.*, 2009; Harriott and Noverr, 2009; Lee *et al.*, 2014). Although different species within biofilms are closely associated and are expected to strongly interact with each other (Elias and Banin, 2012), little is known about how these interactions affect antimicrobial tolerance. Indeed, most previous studies focused on the overall tolerance of multispecies biofilms, without looking at the contributions of individual species. In the limited cases that multispecies composition before and after treatment was determined, the types of interactions and their interdependency with antimicrobial treatment and tolerance were generally not investigated (Harriott and Noverr, 2009; Van der Veen and Abee, 2011; Simões *et al.*, 2009; Whiteley *et al.*, 2001; Leriche *et al.*, 2003; Luppens *et al.*, 2008; Wang *et al.*, 2013; Schwering *et al.*, 2013).

Interactions can be cooperative or competitive in nature. Cooperative interactions for example involve the secretion of enzymes (Rakoff-Nahoum *et al.*, 2016) or metabolic cross-feeding (Harcombe, 2010). Social evolution theory defines a cooperative adaptation in one species as a phenotype that increases the fitness of another species and that evolved at least in part because of this effect (Mitri and Foster, 2013; Foster and Bell, 2012; West *et al.*, 2007). This implies that both species are benefitting from the interaction, since it is difficult to see how an adaptation that helps another species can evolve when it has a fitness cost to the helping species. The *cooperation criterion* therefore states that an observed interaction is only consistent with a cooperative adaptation if the total productivity in co-culture is higher than the sum of the mono-culture productivities (which is the null for no interaction) and both species increase their cellular productivity in co-culture vs. mono-culture (Mitri and Foster, 2013). Higher order cooperative interactions could potentially occur when sets of three, four or more species engage in loops of mutually beneficial interactions. In addition to evolved cooperation, also accidental positive effects can occur (no positive feedback loop), for example when waste products in a focal species can be used as resource by another species. This can result in a commensal interaction, in which there is no fitness effect in one direction, or can happen within a competitive association (*vide infra*).

Whenever one or both species experience a disadvantage in co-culture, competition is dominant. Competition is expected to be favoured when coexisting species have overlapping metabolic niches, are spatially mixed and when cell density is high relative to the available resources (Ghoul and Mitri, 2016). Phenotypes involved in microbial competition can be accidental in nature or have evolved for this purpose. The latter phenotypes are called competitive adaptations and include (i) strategies to take away resources from competitors (exploitative competition), for example by fast-but-wasteful-growth (Pfeiffer *et al.*, 2001), production of nutrient-scavenging molecules (Scholz and Greenberg, 2015), or superior positioning within the niche (Kim *et al.*, 2014), and (ii) strategies to directly fight with competing species (interference competition), for example by production of antimicrobials (Riley and Gordon, 1999) or contact dependent inhibition (Russell *et al.*, 2014; Lories *et al.*, 2017). Competition can be further characterized by comparing the observed productivity in co-culture with the weighted average productivity of the constituent species in mono-culture. This allows to determine to which extent inter-species competition differs from intraspecific competition. As described below, this difference is called the *biodiversity effect* and is constituted of a *selection effect* and *complementarity effect*. Previously, this concept has been frequently applied in plant ecology (Cardinale *et al.*, 2007; Morin *et al.*, 2011; Polley *et al.*, 2003; Spehn *et al.*, 2005; Loreau and Hector, 2001).

Here we combine the *cooperation criterion* and *biodiversity effect* to investigate how inter-species interactions underlie the increased tolerance of multispecies biofilms. We apply our approach to industrially-relevant multispecies biofilm models consisting of combinations of species isolated from contaminating biofilms in breweries. In the brewing industry, undesired biofilms can be associated with spoilage organisms, cause corrosion and reduce process efficiency. Several stages of the brewing process, including the pasteurization, the storage and bottling of the beer, are known to be affected by biofilms and the eradication of these biofilms remains highly challenging (Storgards and Tapani, 2006; Mamvura and Iyuke, 2011; Maifreni *et al.*, 2015). Improved understanding of antimicrobial tolerance is thus a prerequisite for designing more effective brewery sanitation procedures.

Our analysis reveals that a complex interplay between antimicrobial treatment and genetic diversity underlies the commonly-observed increased tolerance of multispecies biofilms. Consistent with previous work in other microbial communities (Foster and Bell, 2012; Rivett *et al.*, 2016; Oliveira *et al.*, 2015), we show that competitive interactions dominate among species within the brewery biofilms. We then show that this dominance of competitive interactions is central to the enhanced antimicrobial tolerance of the multispecies biofilms. Antimicrobial treatment, if incomplete, can reduce the level of competition and therefore cause a subset of species to bloom. The result is a lower percentage inhibition of these species in the multispecies biofilm compared to the mono-culture biofilms, which appears-per definition-as increased tolerance. Complete inhibition of all species in the mixture would avoid this effect. However, our results further indicate that the presence of competing species can also directly enhance the inherent tolerance of microbes to antimicrobial treatment. Antimicrobials that are completely effective against mono-culture biofilms are thus not necessarily effective against the same species in co-culture. Overall this emphasizes that the activity of antimicrobials against multispecies biofilms cannot be predicted based on their effect against mono-cultures.

## MATERIAL AND METHODS

### Sampling

A total of 58 samples were collected from four breweries in Belgium between August and December 2014, before and after cleaning in place (CIP). Sterile cotton swabs (Deltalab) were used for collection of biofilm material from approximately 25 cm^2^ on different surfaces in the bottling plant, filtration room and storage room. After sampling, swabs were submerged in phosphate buffered saline (PBS), which consists of 8.8 g/L NaCl, 1.24 g/L K_2_HPO_4_, and 0.39 g/L KH_2_PO_4_ (pH 7.4). Biofilm material was removed from the swabs by 3× 30 s vortexing and sonication at 45 kHz, 80 W. 900 μl of the PBS solution with the biofilm material was frozen at -80°C in 50% glycerol. 100 μl was plated out in dilution 10° to 10^−8^ on plate count agar (PCA), which is composed of 5 g/L peptone, 2.5 g/L yeast extract, 1 g/L glucose and 15 g/L agar. The plates were incubated at 25 °C for 7 days. The total microbial load was determined by counting the colonies on the plates and determining the CFU/cm^2^.

### Identification of culturable species

Colonies growing on the PCA plates were identified by partial 16S rRNA or ITS gene sequencing. For bacteria, colony PCR using Taq DNA polymerase (Life Technologies) was performed on the 16S rRNA gene, which was targeted by primers BSF8/20 and BSR1541/20 (Cai *et al.*, 2003). The following PCR program was used: 96°C for 6 min, 35 cycles of (i) denaturation at 96°C for 1 min, (ii) annealing at 47.5°C for 1 min, and (iii) elongation at 72°C for 90s, and a final elongation at 72°C for 6 min. Afterwards, the PCR products were loaded on a 1X agarose gel, which was run for 1 hour at 125V and 400A. The band at 1500 bp was cut out and the DNA was extracted by using the GenElute^™^ Gel Extraction Kit (Sigma-Aldrich). Sanger sequencing was performed on the extracted DNA using primer BSF8/20 (GATC Biotech). The resulting sequence was blasted against the NCBI gene database to identify the closest relative of each colony. For yeast identification, the same protocol was followed, using primers ITS1 and ITS4 (White *et al.*, 1990), with an annealing temperature of 60°C and an elongation time of 1 min. The resulting PCR fragment was 330 bp.

### Defined multispecies biofilm models

Defined multispecies biofilm models combined a fixed number of culturable species that were isolated from the same sample. Hereto, each species was grown in liquid PCA culture for 48 hours at 25 °C under shaking conditions. These species were combined with a starting density of 1000 CFU/ml for each species and grown in 1/20 Trypticase Soy Broth in 96 well plates. Similarly, monospecies biofilms were set up with a starting density of 1000 CFU/ml. After incubation for 4 days at 25°C, the amount of living cells in the biofilm formed on the bottom of the 96 well plates was quantified by plate counting. First, the free-living cells were removed from the wells. Second, the biofilm on the bottom of the wells was scraped off in 200 μl PBS and diluted appropriately. Finally, the diluted solutions with biofilm cells were plated out on PCA plates and incubated at 25°C for 2-5 days. Colonies on the plates were counted and the CFU/cm^2^ was determined. In the multispecies biofilms, a distinction was made between the different species based on colony morphology.

### Study of inter-species interactions: cooperation criterion and biodiversity effect

15 different defined multispecies biofilms were grown as described above. To determine if inter-species interactions are cooperative or competitive, the cooperation criterion was applied. This criterion requires that the biofilm growth for all species in co-culture is higher than their respective biofilm growth in mono-culture (Mitri and Foster, 2013).

To further characterize the ecological influences on interactions, the biodiversity effect was calculated according to the formula below, in which 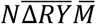 measures the complementarity effect and *N cov*(Δ*RY, M*) measures the selection effect (Loreau and Hector, 2001).

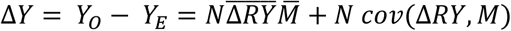

*M*_*i*_ = biofilm growth of species i in mono-culture
*Y*_*o,i*_ = observed biofilm growth of species i in co-culture
*Y*_*o*_ = ∑_*i*_ *Y*_*o,i*_ = total observed biofilm growth in co-culture
*RY*_*E,i*_ = expected relative biofilm growth of species i in co-culture, which is its proportion inoculated
*RY*_*O,i*_ = *Y*_*O,i*_/*M*_*i*_ = observed relative biofilm growth of species i in co-culture
*Y*_*E,i*_ = *RY*_*E,i*_*M*_*i*_ = expected biofilm growth of species i in co-culture
*Y_E_* = ∑_*i*_ *Y*_*E,i*_ = total expected biofilm growth in co-culture
Δ*Y* = *Y*_*o*_ – *Y_E_* = deviation from the total expected biofilm growth in the co-culture (=biodiversity effect)
Δ*RY*_*i*_ = *RY*_*o,i*_ – *RY*_*E,i*_ = deviation from expected relative biofilm growth of species i in co-culture
*N* = number of species in co-culture

The cooperation criterion requires the total multispecies inoculation density to be equal to the sum of the inoculation densities of the mono-culture biofilm. In contrast, the definition of the biodiversity effect imposes that the inoculation density of each species in the multispecies biofilms should be its inoculation density in mono-culture, divided by the number of species that are present in the multispecies biofilm. All multispecies biofilm models in this study were grown in both set-ups and no differences in final growth were observed. Inoculation densities are indeed not expected to have large effects when biofilms are grown to stationary phase (Foster and Bell, 2012).

### Biofilm inhibitor screening

A library of 96 inhibitors that were reported in literature or *in house* developed and that are known to affect biofilm specific processes was composed. Based on commercial availability, low cost and toxicity, 22 compounds were selected for a one-replicate preventive screening against 17 undefined multispecies biofilm, grown directly from the frozen samples. Inhibitors were dissolved in dimethyl sulfoxide (DMSO) at 100 μM and their activity was tested as described in the next paragraph. In a next step, 12 of the 22 inhibitors were selected based on their activity and chemical properties and were screened against 9 of the 17 frozen sample multispecies biofilms using a range of twofold serial diluted concentrations between 800 and 0.4 μM. This allows calculating the BIC50, which is defined as the concentration (μM) of inhibitor needed to inhibit 50% of the biofilm growth.

The undefined multispecies biofilms were set up with a 1000 CFU/ml start inoculum taken directly from the frozen isolated biofilm sample and grown in 1/20 Trypticase Soy Broth on the pegs of the Calgary biofilm device for 4 days at 25 °C. Quantification of biofilm biomass was done by crystal violet staining (Ceri *et al.*, 1999). Briefly, the pegs were first washed in 200 μl PBS and then stained for 30 minutes with 200 μl 0.1% (w/v) crystal violet in an isopropanol/methanol/PBS solution (v/v 1:1:18). Next, the excess stain was washed off the pegs in 200 μl distilled water and the pegs were left to dry for 30 minutes. Finally, the pegs were destained in 200 μl 30 % glacial acetic acid and biofilm matrix was quantified by measuring the OD_570_ of each well using a Synergy MX multimode reader (Biotek, Winooski, VT). The BIC50 was determined for each inhibitor that was tested in a range of concentrations by nonlinear curve fitting (GraphPad Prism software, version 6).

### Multispecies biofilm inhibition by sulfathiazole

Defined mono- and multispecies biofilms were set up in 96 well plates as previously described and at the start of the incubation, sulfathiazole (100 μM in DMSO) or an equal amount of DMSO was added. Biofilms were grown and quantified by plating out as described previously. Inter-species interactions were determined as described above. Tolerance to sulfathiazole was calculated in mono- and multispecies biofilms using the following formulas:

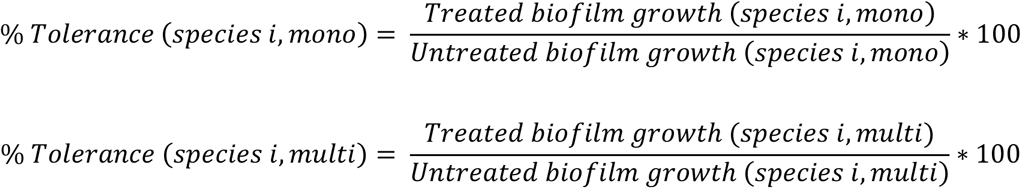

## RESULTS

### Construction of industrially relevant multispecies biofilm models

We started by setting up a series of *in vitro* multispecies biofilm models with relevance for the brewing-industry, that were further used throughout this study. Hereto, 103 biofilm samples isolated from different locations in several breweries were microbiologically characterized. The total bacterial load (CFU/cm^2^) varied between 10^2^ and 10^8^ before cleaning in place (CIP) and between 10^1^ and 10^9^ after CIP. As shown in Figure 1, the microbial contamination after CIP was reduced with less than 75% in 52% of the samples and was even increased in 24% of the samples, indicating that CIP is insufficient and that improved antimicrobial treatments are highly needed.

**Figure 1:**
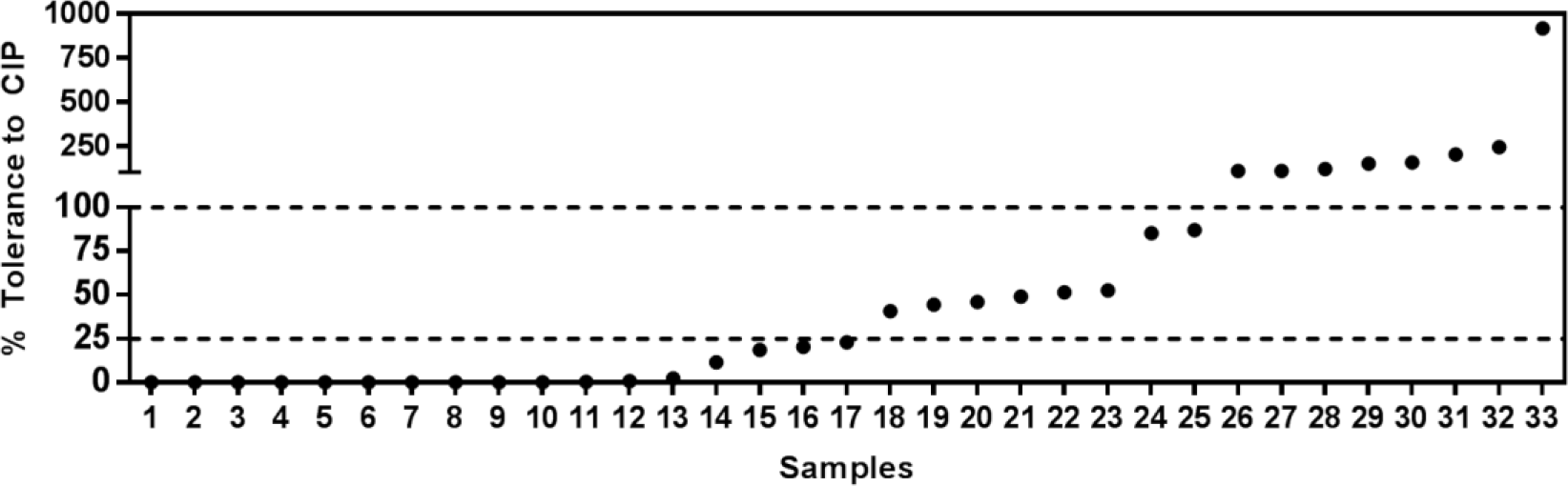
% tolerance to CIP for 33 biofilms sampled from different locations in different breweries.

The genera of the closest known relatives of the culturable microbes were determined by partial 16S rRNA gene sequencing to characterize the microbial diversity (Table S1). The biofilm samples were mainly composed of *Pseudomonas* and *Raoultella* ssp. and also two beer spoiling organisms, *Pediococcus* and *Lactococcus*, were identified. Multispecies biofilm models were then constructed by combining species isolated from the same sample. Seventeen ‘undefined’ biofilm models were set up by directly inoculating part of the frozen isolated biofilm samples. These models were used to screen for broad-spectrum biofilm inhibitors. Because these biofilms likely contain unculturable species, for the study of inter-species interactions an additional 12 ‘defined’ multispecies biofilm models were constructed by inoculating equal ratios of 3 to 6 well-identified, culturable species (originating from 8 samples taken before CIP and 4 samples taken after CIP). Several biofilms containing *Pseudomonas* and *Raoultella* spp. were included.

### Screening of biofilm inhibitors

To study the tolerance of multispecies brewery biofilms, we sought to use a broad-spectrum antimicrobial with a large potential for application against brewery biofilms. Hereto, a library of 22 biofilm inhibitors -with previously reported activity against mono-culture biofilms- was composed that target biofilm specific processes such as adhesion (Opperman *et al.*, 2009), dispersion (Barraud *et al.*, 2006), EPS-production (Nithya *et al.*, 2011) and several others (Lynch and Abbanat, 2010). After an initial screening against 17 undefined multispecies biofilm models using a fixed concentration of 100 (data not shown), we selected 12 inhibitors, which were tested more thoroughly using multiple concentrations. Specifically, we performed a preventive screening against 9 undefined multispecies biofilm models directly grown from the frozen brewery biofilm samples. Crystal violet staining was used to measure the amount of biofilm formed and the 50% inhibitory concentrations (BIC50) were calculated for each biofilm model (Figure 2). Sulfathiazole was found to have the broadest activity-spectrum against the brewery biofilms and was therefore selected for further study. This inhibitor has been described previously to interfere with c-di-GMP biosynthesis in *E. coli* biofilms (Antoniani *et al.*, 2010). C-di-GMP has been reported to play a crucial role in biofilm formation by a wide range of bacterial species, which might explain the broad-spectrum activity of this compound (Cotter and Stibitz, 2007; Hengge, 2009; Romling *et al.*, 2013).

**Figure 2:**
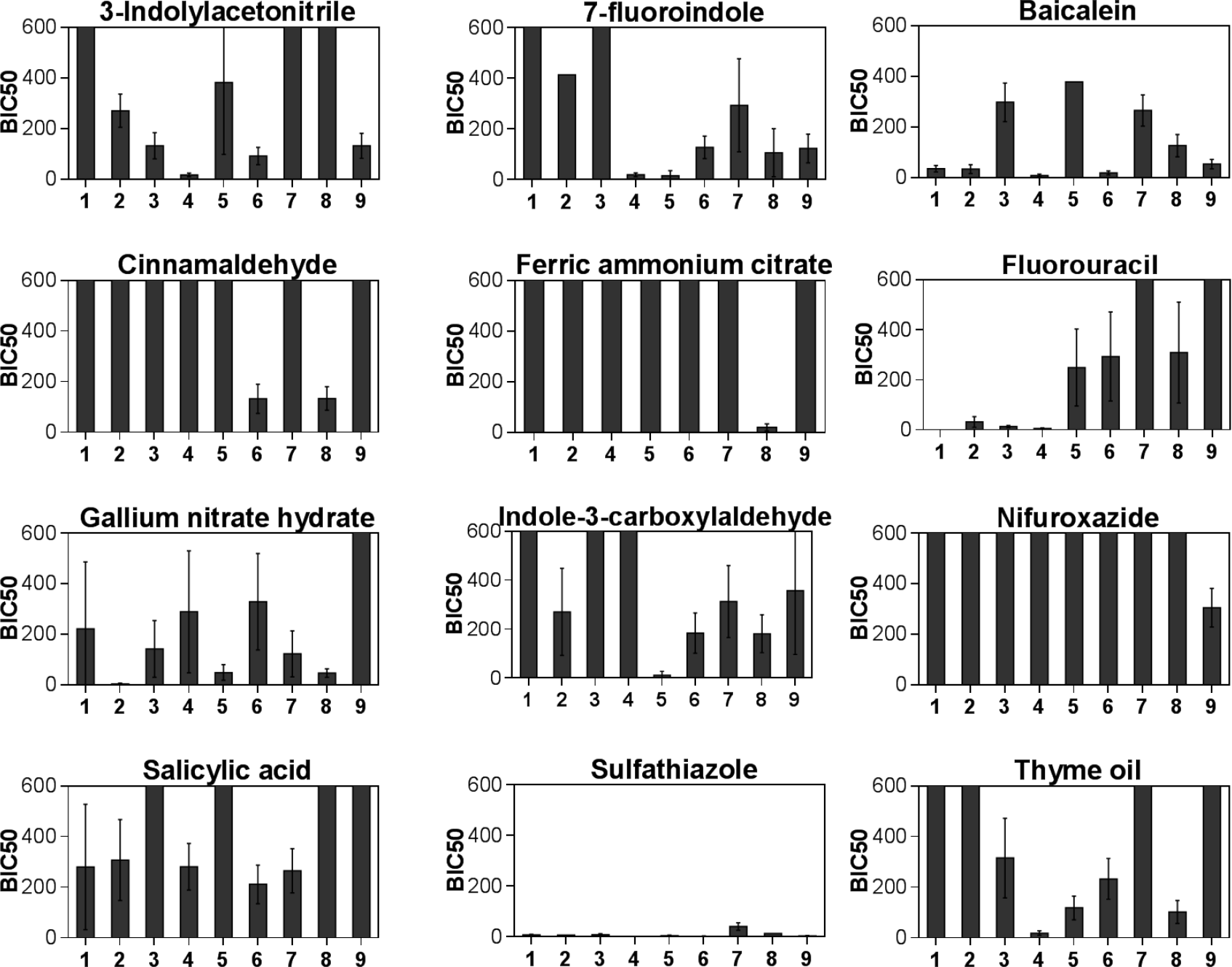
BIC50 values of 12 biofilm inhibitors against 9 undefined biofilms (shown as mean with standard deviation of three biological repeats). BIC50 is defined as the concentration (μM) of inhibitor needed to prevent biofilm growth with 50%. Compounds with BIC50 values over 600 μM are considered ineffective and are not shown.

### Effect of inter-species interactions on biofilm growth and composition

We first aimed to determine the role of inter-species interactions in multispecies biofilms, irrespective of antimicrobial treatment. Hereto, we performed a systematic classification of the interactions in 12 defined biofilm models, each consisting of 3 to 6 culturable species, by using two complementary approaches: (i) *cooperation criterion* and (ii) *biodiversity effect*.

The *cooperation criterion* was used to classify interactions as cooperative or competitive. For all biofilm models, the number of biofilm cells of each species (CFU/cm^2^) in mono-culture was compared to the cell count of each species in co-culture (Figure 3). Most species performed worse in co-culture than in mono-culture indicating that competitive interactions are dominant. The increased cellular productivity observed for a subset of the species in few of the multispecies biofilms (e.g. for 3 out of 4 species in model 5) could be due to exploitation of the remaining, suppressed species, however, cooperation between these species cannot be ruled out. In summary, all models were characterized by competitive interactions that cause some or all species to perform worse than in mono-culture.

**Figure 3:**
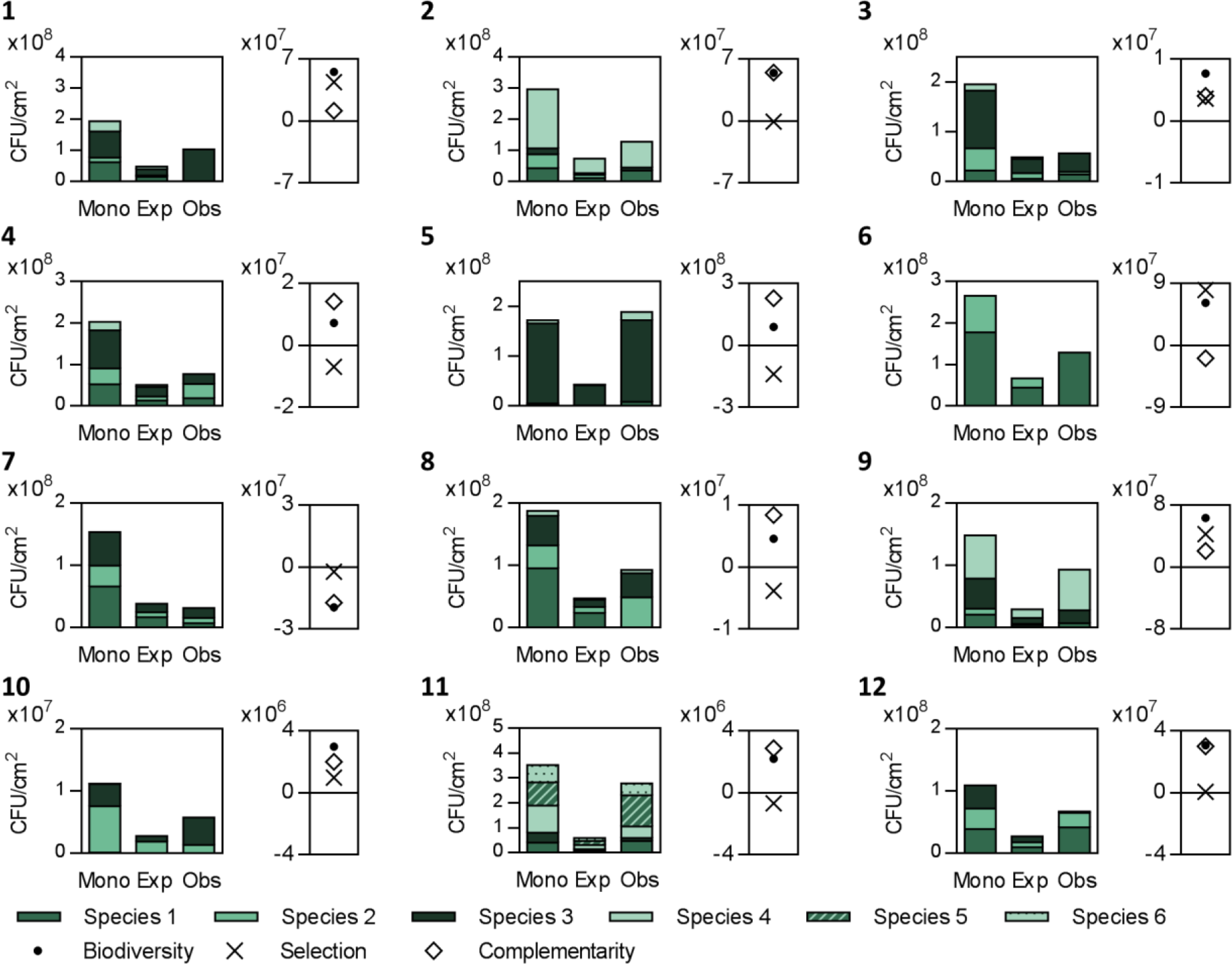
Mono-culture growth (Mono), expected (Exp) and observed (Obs) multispecies composition and the biodiversity, selection and complementarity effect for 1 representative repeat of 12 defined multispecies biofilm models.

To further characterize competition, the *biodiversity effect* was measured (Loreau and Hector, 2001). When inter-species competition is equal to intra-specific competition, the observed productivity in co-culture is expected to be equal to the average productivity of the constituent species in mono-culture, weighted by the inoculation frequencies. The biodiversity effect is defined as the difference between the observed and expected multispecies biofilm productivity and is thus a measure for the extent to which inter-species interactions deviate from intra-specific interactions. The observed productivity of the 12 model biofilms (CFU/cm^2^) was compared to the expected biofilm growth. In the majority of the multispecies biofilm models (75%), the total amount of biofilm formed was higher than expected, as indicated by a positive biodiversity effect, while the remaining 25% of the cases were characterized by a negative biodiversity effect.

A positive biodiversity effect can either be caused by selection of the best biofilm former or by a (partial) niche separation alleviating competition; conversely a negative biodiversity effect can be caused by selection of the worse biofilm former or by strong interference competition. To distinguish between both possibilities, Loreau & Hector (2001) partitioned the biodiversity effect into a selection and complementarity effect (Material and Methods). Selection occurs when the extent to which the relative productivity in co-culture vs. mono-culture deviates from expected is non-randomly related to the productivity in mono-culture and is measured by a covariance function. Positive selection is indicative of the dominance of the best mono-culture biofilm formers and occurred in 33,3% of the multispecies biofilm models. Negative selection suggests the opposite and appeared in the remaining 67,7% of the multispecies biofilm models. If only selection effects take place, the total relative productivity (sum of relative productivities of all species) is 1, meaning that an increase in productivity in one species is compensated by a decrease in productivity of another species. However, if the total relative productivity is higher or lower than 1 over- or underyielding occurs, which is defined as the complementarity effect. This effect measures whether the relative amount of biofilm formed in co-culture vs. mono-culture is on average higher or lower than expected based on the initial relative abundance and biofilm growth in mono-culture and is thus also a measure for the strength of competition. Complementarity is positive if some degree of niche separation occurs, for example if two species can grow on different resources or if one species is able to use a waste product of another species as a resource. Consequently, the strength of competition decreases and the productivity increases due to a more optimal use of the available niches. Positive complementarity was observed in 91,7% of the multispecies biofilm models. On the other hand, negative complementarity effects occurred in the remaining 8,3% of the multispecies biofilm models and indicate the occurrence of strong chemical or physical interference competition (Fox, 2005; Turnbull *et al.*, 2013; Loreau, 2000).

The combination of complementarity and selection effects then gives an indication as to which ecological processes are the cause of the total positive or negative biodiversity effect. In our multispecies biofilm models positive biodiversity (75%), could be explained by resource partitioning or facilitation between the different species for 66,7% of the biofilms (only positive complementarity), by dominance of the best biofilm formers for 11,1% of the biofilms (only positive selection) or by a combination of both positive complementarity and selection for 22,2% of the biofilms. Conversely, negative biodiversity effects (25%), were caused by exploitation or interference competition for 33,3% of the biofilms (only negative complementary), by dominance of poor biofilm formers for 33,3% of the biofilms (only negative selection) or by a combination of negative complementarity and selection for 33,3% of the biofilms (Loreau and de Mazancourt, 2013). Overall, the mainly positive complementarity effects indicate that the competitive interactions in the multispecies biofilm models are in most cases alleviated by partial niche separation.

### Link between reduced competition and antimicrobial tolerance in multispecies biofilms

The results above show that competitive inter-species interactions, although in general alleviated by partial niche separation, strongly influence the productivity of each species in the multispecies biofilms. In a next step, we sought to investigate the interplay of these competitive interactions with antimicrobial treatment and their effect on antimicrobial tolerance. Hereto, sulfathiazole was added preventively to three multispecies biofilm models (Table 1). Tolerance to sulfathiazole is defined as the ratio between the amount of biofilm formed in the presence and absence of treatment and was determined for each species in mono- and co-culture conditions (Figure 4-6). In all three models the tolerance of each species was equal or higher in the multispecies biofilm than in the mono-culture. The result is an overall increase in tolerance in each multispecies biofilm, which can be seen by comparing the expected and observed amount of biofilm after treatment. Here the expected amount is calculated based on the composition before treatment and the percentage of reduction of each species in mono-culture. These results are in line with the increased tolerance generally observed in multispecies biofilms (Mozina *et al.*, 2013; Burmølle *et al.*, 2014). Specifically, our results are consistent with a previous study on sulfathiazole treatment, in which multispecies biofilms isolated from cooling water systems were found to be more tolerant compared to their mono-culture counterparts (Shakeri *et al.*, 2007).

**Table 1:** Closest known relative genus for each of the species present in the three multispecies biofilm models that were used to study the inhibition by sulfathiazole

|  | Species 1 | Species 2 | Species 3 | Species 4 |
| --- | --- | --- | --- | --- |
| Model 1 | *Epilithonimonas* | *Aeromonas* |  |  |
| Model 2 | *Pseudomonas* | *Pseudoclavibacter* | *Raoultella* | *Serratia* |
| Model 3 | *Pseudomonas* | *Raoultella* |  |  |

**Figure 4:**
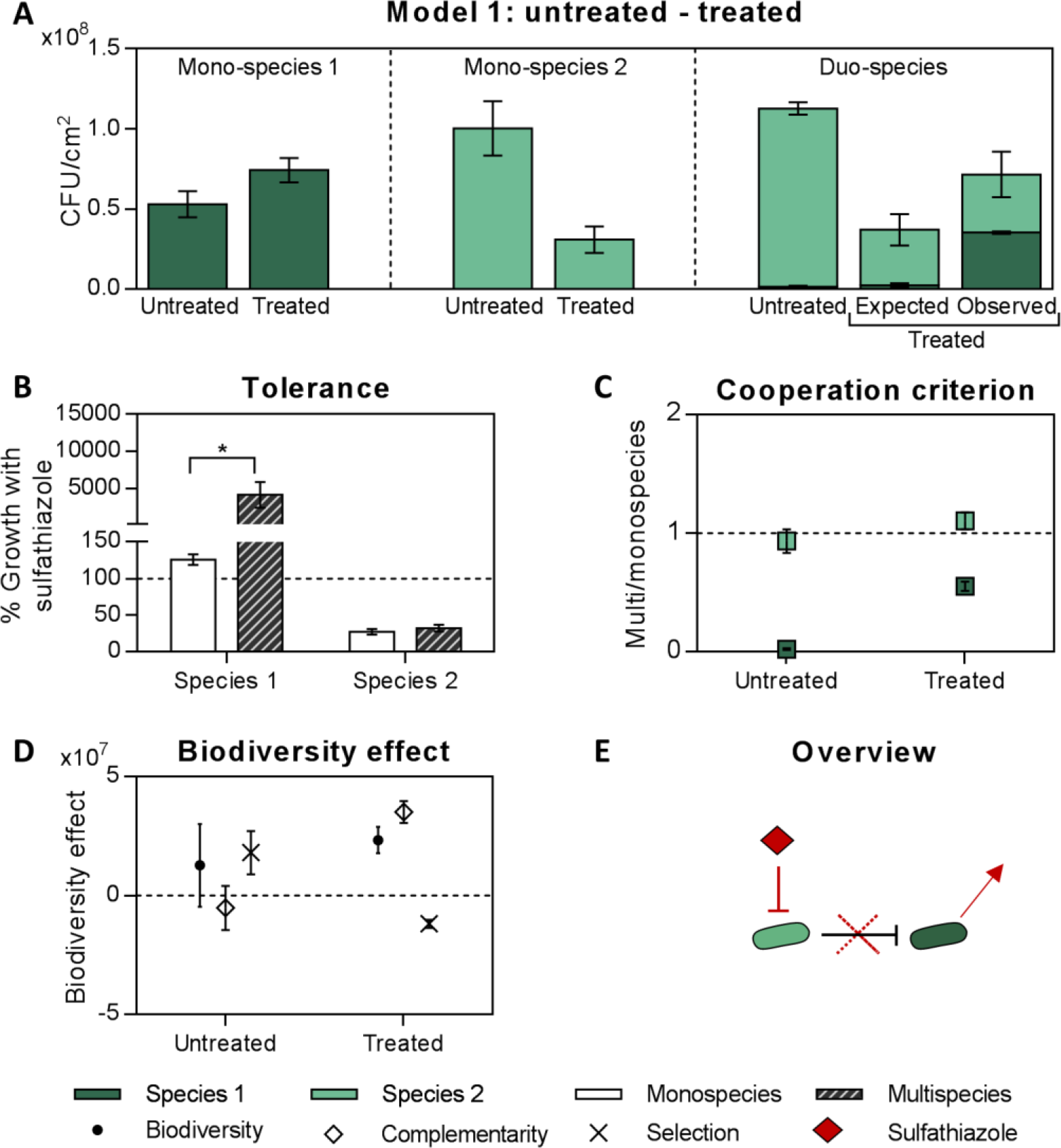
Model 1: **A**: Number of cells of each species in mono- and co-culture biofilms, grown in the absence and presence of sulfathiazole treatment: suppressed species 1 is able to grow after treatment. **B**: Tolerance = ratio between the number of biofilm cells with and without sulfathiazole treatment, determined for each species in mono- and co-culture conditions. For each species the tolerance is equal or higher within the co-culture biofilm. Significant differences were examined using a two-way anova and Bonferroni correction (* P<0.05) **C**: Cooperation: both in the absence and presence of treatment the criterion for cooperation is not met. **D**: Biodiversity effects in the absence and presence of treatment of the duo-species biofilm: dominating positive selection is replaced by positive complementarity. **E**: Overview: inhibition of species 2 leads to a reduction in the competitive interactions against species 1, which allows species 1 to bloom. Results show the average of 3 biological repeats, except for A, which shows the average of 3 technical repeats of one representative biological repeat.

Analyzing biofilm compositions before and after treatment revealed that the above described dominance of competitive interactions in untreated biofilms is central to the observed enhanced tolerance to antimicrobial treatment. In two out of three biofilm models we found that antimicrobial treatment reduced the level of competition and therefore caused a subset of species to bloom. The result was a lower percentage inhibition of these species in the multispecies biofilm compared to the mono-culture biofilms, which -per definition- appears as increased tolerance.

In duo-species model 1 (Figure 4), species 2 is sensitive to the inhibitor both in the mono-and co-culture biofilm (4 A&B). However, species 1, which is insensitive to the inhibitor in mono-culture, shows a 50-fold increase in growth upon addition of the inhibitor in the duo-species biofilm (4 A&B), resulting in an overall higher tolerance of the duo-species biofilm (4A). In the untreated duo-species biofilm, species 1 is strongly suppressed by species 2 as reflected in the strong competition (4C), large positive selection effect (4D) and negative complementarity (4D). The increased growth of species 1 upon treatment is therefore consistent with an abrogation of the competitive interactions of sensitive species 2 against species 1, which then blooms and shows a net increase in antimicrobial tolerance. This is reflected in a reduced competition (4C), associated with a positive complementarity (4D) in the treated biofilm. In summary, inhibition of the best competitor results in a bloom of the worse competitor and overall increased tolerance.

A similar mechanism plays in tetra-species model 2 (Figure 5). Three out of four species are completely inhibited both in the mono-and multispecies biofilm (5 A&B). Species 3, however, which is insensitive to the inhibitor in the monospecies biofilm, shows a 1.5-fold increase in growth upon addition of the inhibitor in the multispecies biofilm (5 A&B), resulting in an overall increase in tolerance of the mixed species biofilm (5A). Species 3 experiences competition by the other species in the untreated multispecies biofilm (5 C&D), explaining why inhibition of these other species increases the growth-and tolerance-of species 3 in the treated biofilm. Since there is only one species left after treatment, competition (5C) and biodiversity effect (5D) are zero.

**Figure 5:**
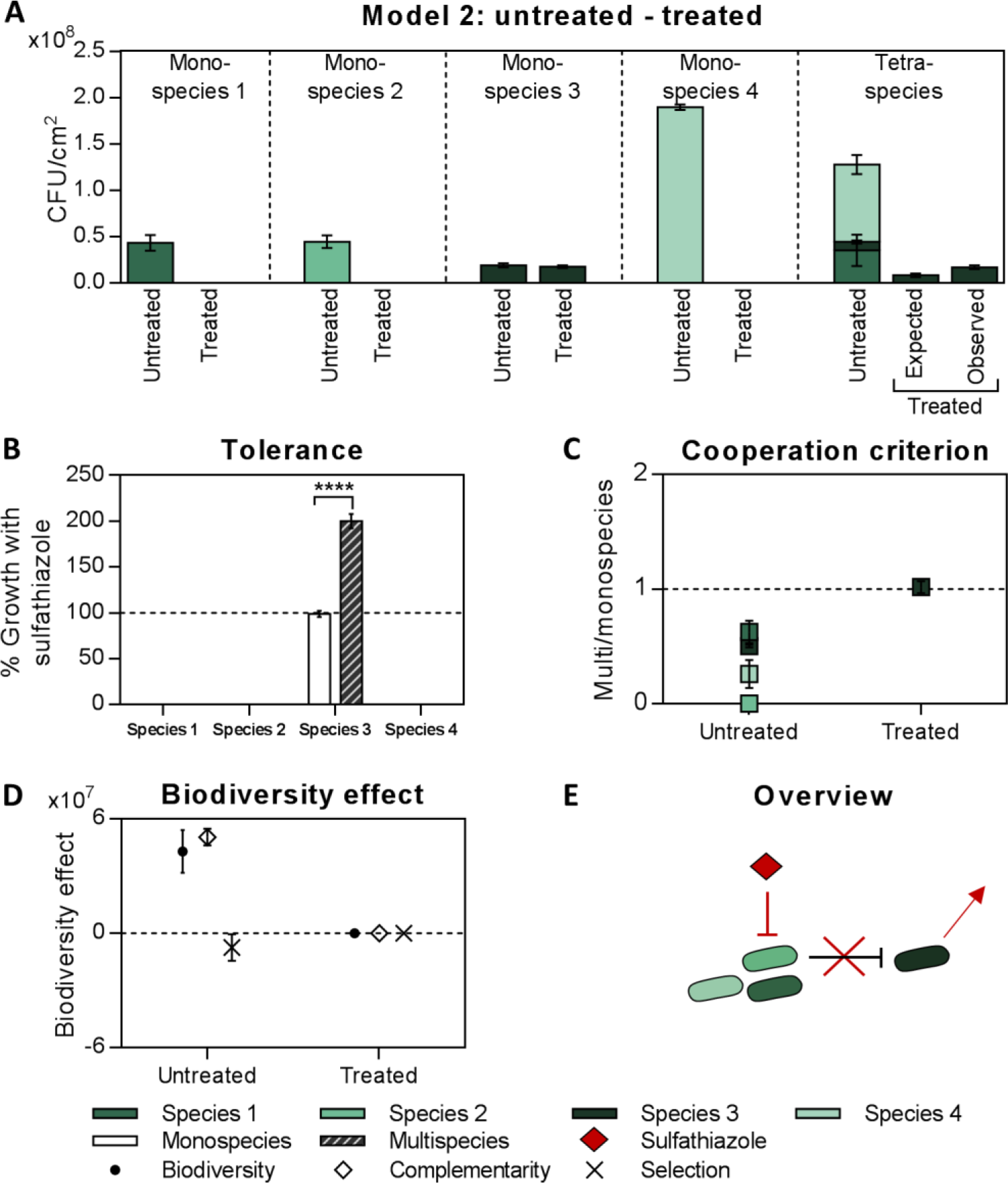
Model 2: **A**: Number of cells of each species in mono- and co-culture biofilms, grown in the absence and presence of sulfathiazole treatment: suppressed species 3 shows an increased growth upon treatment. **B**: Tolerance = ratio between the number of biofilm cells with and without sulfathiazole treatment, determined for each species in mono- and co-culture conditions. Species 3 shows an increased the tolerance within the multispecies biofilm, while species 1, 2 and 4 are completely inhibited in mono- and co-culture. Significant differences were examined using a two-way anova and Bonferroni correction (**** P<0.0001). **C**: Cooperation: both in the absence and presence of treatment the criterion for cooperation is not met. **D**: Biodiversity effects in the absence and presence of treatment of the multispecies biofilm: negative complementarity becomes positive. **E**: Overview: complete inhibition of species 1, 2 and 4 leads to the abrogation of competitive interactions against species 3, which allows species 3 to bloom. Results show the average of 3 biological repeats, except A, which shows the average of 3 technical repeats of one representative biological repeat.

It should be noted that this mechanism of ‘increased tolerance due to reduced competition’ does not involve an increase in absolute cell numbers of the different species in co-culture compared to mono-culture, nor an expression of specific tolerance phenotypes. Nevertheless, the proposed mechanism is of significance. Indeed, similar to our study, antimicrobial tolerance in previous studies was generally measured by calculating the reduction in cell numbers before and after treatment, not by directly comparing the absolute cell numbers between co- and mono-culture conditions (Chorianopoulos *et al.*, 2008; Van der Veen & Abee, 2011; Kostaki *et al.*, 2012; Giaouris *et al.*, 2013; Wang *et al.*, 2013). Therefore, the increased tolerance observed in these studies might as well be explained by decreased competition and should not necessarily be accompanied by any changes in specific tolerance phenotypes.

### Direct effect of competitors on antimicrobial tolerance

The findings above indicate that incomplete antimicrobial treatment of multispecies biofilms can reduce the levels of competition and therefore cause a subset of species to bloom, which ultimately results in increased antimicrobial tolerance. Complete inhibition of all species in the mixture would solve this problem. However, our analysis of duo-species model 3 (Figure 6) indicates that the presence of competing species can also directly enhance the inherent tolerance of other species by driving specific tolerance phenotypes. This means that antimicrobials that are completely effective against mono-culture biofilms are not necessarily effective against the same species in co-culture and thus precludes any prediction on multispecies tolerance.

**Figure 6:**
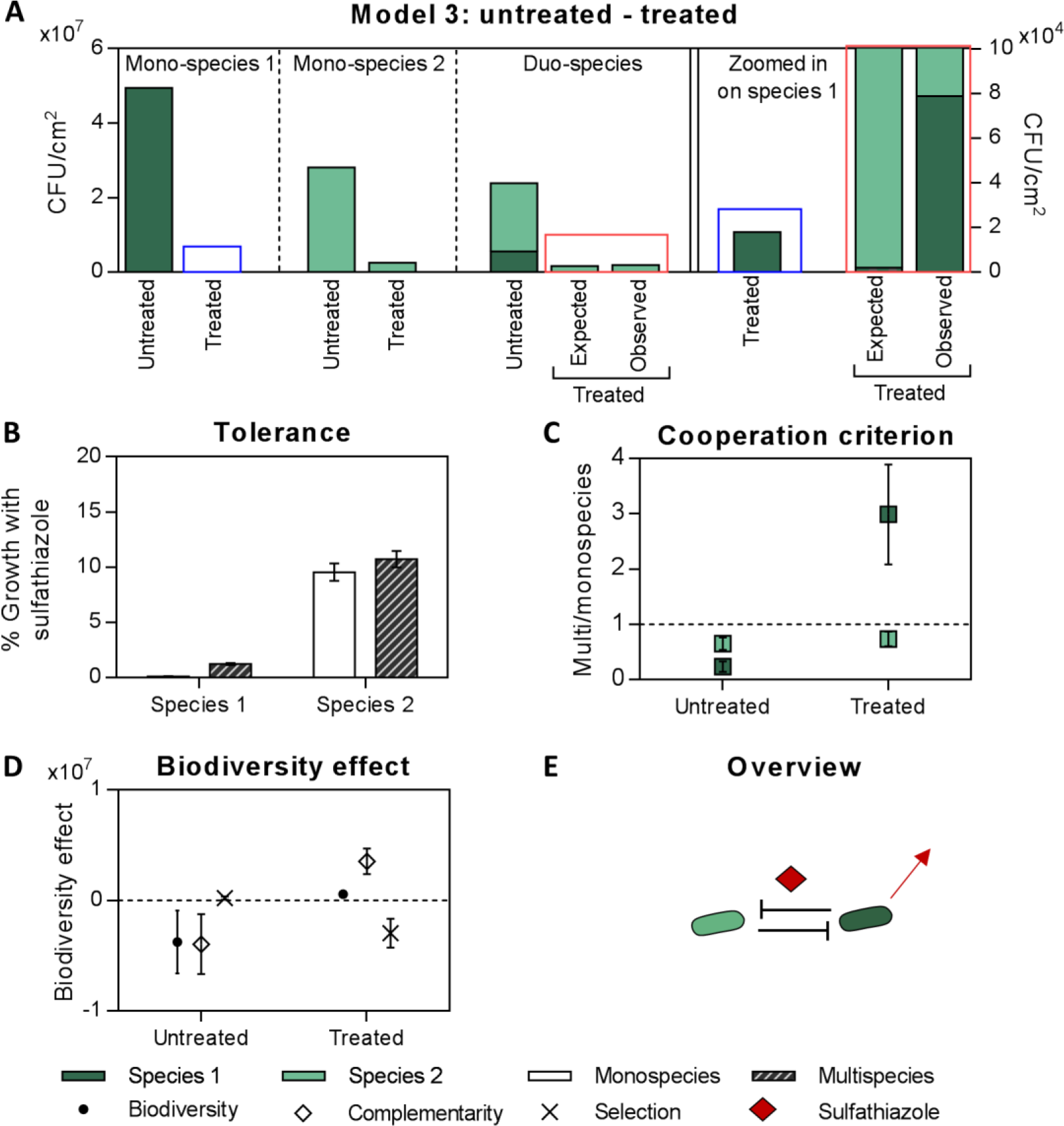
Model 3: **A**: Number of cells of each species in mono- and co-culture biofilms, grown in the absence and presence of sulfathiazole treatment. The right part of the graph zooms in on the amount of biofilm formed by species 1 in treated mono- and duo-culture: after treatment the growth of species 1 in duo-culture exceeds its growth in mono-culture. **B**: Tolerance = ratio between the number of biofilm cells with and without sulfathiazole treatment, determined for each species in mono- and co-culture conditions. Species 1 shows a higher tolerance within the co-culture biofilm, while there is no difference for species 2. **C**: Cooperation: after treatment species 1 grows better in co-culture than in mono-culture, while there is no difference for species 2. This is consistent with commensalism. **D**: Biodiversity effects in the absence and presence of treatment of the co-culture biofilm: negative complementarity before treatment becomes positive. **E**: Overview:, species 1 becomes more tolerant in the presence of competing species 2. The growth of species 1 in the treated co-culture biofilm even exceeds its mono-culture growth, suggesting induction of specific tolerance phenotypes Results show the average of 3 biological repeats, except A, which shows one representative biological.

In this duo-species model (Figure 6), both species respond to sulfathiazole treatment in the mono- and co-culture biofilms (6 A&B). However, species 1 shows a 11,1-fold reduction in sensitivity in co-culture, resulting in an overall increased tolerance of the co-culture biofilm (6A). In contrast to the previous model systems, this tolerance of species 1 is associated with an increase in cell number above the mono-culture levels (6A, right panel). These results cannot be explained by a decrease in competition alone (6C-D),and should be attributed to the presence of specific tolerance phenotypes within the multispecies biofilm. These could either be related to a protective effect of species 2 on species 1 or to a direct change in tolerance phenotype of species 1 as a response to species 2.

## DISCUSSION

Functional properties like antimicrobial tolerance strongly differ between multispecies and monospecies biofilm communities (Burmølle *et al.*, 2014; Røder *et al.*, 2016). Although inter-species interactions are expected to be both intense and important within dense communities (Elias and Banin, 2012), little is known about how they affect antimicrobial tolerance. Previous work either focused on microbial interactions in untreated biofilms (Tan *et al.*, 2016; Røder *et al.*, 2016; Ghoul and Mitri, 2016) or on the overall tolerance of multispecies biofilms, without taking contributions of individual species into account (Adam *et al.*, 2002; Burmølle *et al.*, 2006; Baffone *et al.*, 2011; Simões *et al.*, 2010; Lopes *et al.*, 2012). We have bridged the gap and shown that a complex interplay between antimicrobial treatment and inter-species interactions underlies the commonly-observed increased tolerance of multispecies biofilms. We have shown that competitive interactions dominate within industrially relevant multispecies biofilm models and that antimicrobial treatment, if incomplete, can reduce the level of competition and therefore cause subsets of species to bloom, ultimately leading to enhanced overall tolerance. In addition, we have shown that the presence of competitors can also directly enhance the inherent tolerance to antimicrobials by driving specific tolerance phenotypes. Overall, our results emphasize that the increasingly-recognized dominance of competition in multispecies biofilms is central to the enhanced antimicrobial tolerance and that antimicrobial activities against mono-culture biofilms cannot predict efficacy against multispecies biofilms.

Our data indicate that competitive interactions dominate among species within brewery biofilms, although inter-species competition is generally weaker than intraspecific competition. These data fit with a growing body of recent theoretic and experimental work motivating that competition, not cooperation, dominates interactions among microbial species. The genotypic view of social interactions predicts a low chance of evolution of cooperation between species, because this requires both a high within-genotype relatedness and sufficient niche separation to reduce ecological competition. Increased niche separation, however, often implies a decreased exchange of resources, which counteracts interactions, and further complicates the evolution of cooperation (Mitri and Foster, 2013). These predictions are confirmed by recent systematic screenings of inter-species interactions based on the cooperation criterion (Foster and Bell, 2012; Rivett *et al.*, 2016; Fiegna *et al.*, 2015). Also in these studies inter-species competition was found to be weaker than intraspecific competition (Foster and Bell, 2012; Rivett *et al.*, 2016; Oliveira *et al.*, 2015; Fiegna *et al.*, 2015). Moreover, Rivett *et al.* (2016) showed that initially strong competitive interactions can weaken over time by divergence in resource use and increased niche complementarity. It should be noted that a number of studies did report a prevalence of positive interactions, however, these studies made use of alternative definitions. In a recent study, synergistic interactions were defined as the total amount of multispecies biofilm being higher than the sum of all mono-cultures and synergy was observed in 13% of the biofilms (Madsen *et al.*, 2016). This definition is similar to the cooperation criterion, but since the effect of growth in co-culture on the individual species is not included, the presence of cooperative interactions cannot be confirmed. In earlier studies, synergy required the total amount of multispecies biofilm to be higher than that of the best mono-culture biofilm former and synergistic interactions were reported in respectively 11%, 63% and 30% of the biofilms (Burmølle *et al.*, 2007; Ren *et al.*, 2015; Røder *et al.*, 2015). Also here information on composition of the multispecies biofilm is needed to determine whether the described synergistic interactions are competitive or cooperative. It can however be deduced that these synergistic interactions are associated with a positive biodiversity effect, since both definitions imply the total amount of multispecies biofilm to be higher than the weighted average of the mono-cultures. Notably, this positive biodiversity effect does imply niche complementarity, but can also partly be caused by positive selection effects. In conclusion, the importance of competition among species over cooperation is increasingly recognized and our data are consistent with this. However, an important note is that all studies described above, including ours, are based on culturable species, which might exclude species that are only able to grow in the presence of other species. Therefore, the prevalence of cooperation might be underestimated (Foster and Bell, 2012; Røder *et al.*, 2016).

The enhanced overall antimicrobial tolerance against sulfathiazole that we observed for each multispecies biofilm model compared to the mono-culture biofilms is consistent with the enhanced resistance found in the majority of multispecies biofilm studies (Baffone *et al.*, 2011; Simões *et al.*, 2010; Shakeri *et al.*, 2007; Kumar and Peng, 2015; Jagmann *et al.*, 2015; Adam *et al.*, 2002; Lopes *et al.*, 2012; Leriche *et al.*, 2003; Whiteley *et al.*, 2001; Luppens *et al.*, 2008; Wang *et al.*, 2013; Schwering *et al.*, 2013; Van der Veen and Abee, 2011; Simões *et al.*, 2009; Harriott and Noverr, 2009; Lee *et al.*, 2014; Hoffman *et al.*, 2006). In most of these studies the enhanced tolerance was attributed to protective effects of the species on each other, however, generally without unraveling the mechanism of tolerance. In contrast, a minority of studies did not observe an effect of multispecies conditions on antimicrobial tolerance (Gkana *et al.*, 2017) or did even measure a decrease in tolerance in multispecies conditions (Lindsay *et al.*, 2002; Chorianopoulos *et al.*, 2008; Kart *et al.*, 2014; Yassin *et al.*, 2016; Feldman *et al.*, 2016).

Our data indicate that the commonly-observed enhanced antimicrobial tolerance of multispecies biofilms is associated with a reduction in the level of competition upon treatment, causing a subset of species to bloom. The dominance of competition among species over cooperation in untreated biofilms is therefore central to the enhanced antimicrobial tolerance. Indeed, incomplete inhibition of a network of cooperating species is expected, not to promote, but to pull down the remaining species because of abrogation of positive feedback loops, as is motivated by recent ecological network studies (Coyte *et al.*, 2015). This would reduce, not increase, the overall tolerance of the multispecies biofilm (Feldman *et al.*, 2016). Our models only provide examples of multispecies biofilms in which specific species strongly suppress other species. Inhibition of the stronger competitors consequently reduces the competition that is experienced by the suppressed species and leads to an increased tolerance of the weaker competitors. However, the idea that antimicrobial tolerance in multispecies biofilms is connected to a reduction in competition should not be limited to this situation, as one can easily imagine that antimicrobial treatment can also reduce competition between equal competitors. For example, in the case of equally competing species that only produce their toxins when the population density of the other species is sufficiently high (Cornforth and Foster, 2013), a reduction of the population size by antimicrobial treatment would interfere with toxin production, reduce competition and ultimately lead to increased antimicrobial tolerance compared to mono-culture.

Based on the commonly found prevalence of competitive interactions within multispecies biofilms, it is expected that reduction in competition might often be the cause of increased tolerance. However, little is known about this because previous work mainly focused on characterizing the antimicrobial tolerance of mono- and multispecies biofilms, without explicitly classifying the changes in inter-species interactions before and after treatment. In a number of studies, only the overall activity against the multispecies biofilm and the activity against the mono-cultures was measured, while information on individual species in co-culture is essential to understand the inter-species interactions (Adam *et al.*, 2002; Burmølle *et al.*, 2006; Baffone *et al.*, 2011; Simões *et al.*, 2010; Lopes *et al.*, 2012). Similarly, only determining the inhibition of each species in the co-culture without looking at the effects in mono-culture (Norwood and Gilmour, 2000; Hill *et al.*, 2010; DeLeon *et al.*, 2014; Sun *et al.*, 2008; Feldman *et al.*, 2016) or only focusing on specific species within the multispecies biofilm (Kumar and Peng, 2015; Jagmann *et al.*, 2015; Shakeri *et al.*, 2007) does not allow to study all changes in inter-species interactions. Nevertheless, a few studies have been conducted in which the tolerance of each species was examined individually, both under mono- and co-culture biofilm conditions (Harriott and Noverr, 2009; Van der Veen and Abee, 2011; Simões *et al.*, 2009; Whiteley *et al.*, 2001; Leriche *et al.*, 2003; Luppens *et al.*, 2008; Wang *et al.*, 2013; Schwering *et al.*, 2013; Elvers *et al.*, 2002). While the obtained data would allow to perform a detailed analysis of the changes in inter-species interactions as proposed in this paper, this analysis is generally missing and the representation of the data in most cases did not allow us to interpret the data a posteriori. Nevertheless, one study on tolerance of a 7-species biofilm provided sufficient data and is consistent with our mechanism of ‘increased tolerance due to reduced competition’ (Elvers *et al.*, 2002). Some of the bacterial species experienced a reduced growth due to competition in the untreated multispecies biofilm, while antimicrobial treatment restored their growth in the multispecies biofilm to the level of the untreated monoculture biofilms. In contrast, but also consistent with our rationale, a reduction in antimicrobial tolerance under multispecies conditions has been explicitly associated with a reduction of (probably rare) cooperative inter-species interactions (Feldman *et al.*, 2016).

In our final model, we found that the presence of competitors can also directly enhance the inherent tolerance of other species by driving specific tolerance phenotypes. This could either be attributed to (i) protective effects of specific species on other species or to (ii) direct changes in tolerance phenotypes of specific species as a response to competitors. A previously described example of a protective effect occurs between competing *Pseudomonas aeruginosa* and *Staphylococcus aureus* species (Hoffman *et al.* 2006). Respiration of *S. aureus* was found to be inhibited by a competitive interaction involving the exoproduct 4-hydroxy-2-heptylquinoline-N-oxide of *P. aeruginosa*. As a consequence, aminoglycoside antibiotics were no longer taken up by *S. aureus* cells and their tolerance to these antibiotics increased. Additionally, the presence of *P. aeruginosa* on a long term increased the production of highly resistant small-colony variants of *S. aureus*, which further improved the antimicrobial tolerance of *S. aureus*. In addition, it is becoming increasingly clear that bacteria can also directly sense the presence of competitors and respond appropriately (i.e. ‘competition sensing’) (Cornforth and Foster, 2013). Recent studies indicate that these responses can include upregulated biofilm formation (Oliveira *et al.*, 2015), increased antibiotics or toxin production (Le Roux *et al.*, 2015; Abrudan *et al.*, 2015; Rosenberg *et al.*, 2016), altered secretion of specific secondary metabolites (Traxler *et al.*, 2013), but also increased antibiotic tolerance (Abrudan *et al.*, 2015; Roberfroid *et al.*, personal communication).

In conclusion, due to the their commonly observed increased antimicrobial tolerance, multispecies biofilms remain challenging to eradicate. Accordingly, we found multispecies biofilms to be a serious problem in breweries, as emphasized by the high microbial load of the isolated biofilm samples, both before and after CIP. An increased knowledge of the properties of these multispecies biofilms may aid to improve their control. Our study demonstrates that competitive inter-species interactions dominate within multispecies biofilms and have a strong influence on the outcome of antimicrobial treatment. Specifically, we found that strongly suppressed species can bloom after inhibition of superior competitors by antimicrobial treatment, which results in increased tolerance. To avoid such unwanted effects of changing inter-species interactions, it would be useful to develop combination therapies that completely inhibit all species. Nevertheless, we also observed that the presence of competitors can increase the intrinsic tolerance of species by driving specific tolerance phenotypes. This means that antimicrobials that are completely effective against mono-culture biofilms are not necessarily effective against the same species in co-culture. Our study therefore underlines the need to further investigate and interfere with the mechanisms behind these specific tolerance phenotypes.

## ACKNOWLEDGEMENTS

We would like to thank Stijn Robijns, Sandra Van Puyvelde and Bram Lories for their valuable comments and for their assistance in sampling of brewery biofilms. We thank K. Deflem and D. De Coster for experimental assistance. This work was supported by the KU Leuven Research Fund (STG/16/022), by the Institute for the Promotion of Innovation through Science and Technology in Flanders under grant IWT-SBO 120050 (NEMOA) and by FWO-Vlaanderen (W0.009.16N). IP is a research assistant of the IWT-Vlaanderen (SB/131721). HS acknowledges the receipt of a postdoctoral fellowship from FWO-Vlaanderen (PDO/11).

